# Variant Set Enrichment: An R package to Identify Dis-ease-Associated Functional Genomic Regions

**DOI:** 10.1101/077990

**Authors:** Musaddeque Ahmed, Richard C. Sallari, Haiyang Guo, Jason H. Moore, Housheng Hansen He, Mathieu Lupien

## Abstract

**Summary:** Genetic predispositions to diseases populate the noncoding regions of the human genome. Delineating their functional basis can inform on the mechanisms contributing to disease development. However, this remains a challenge due to the poor characterization of the noncoding genome. Variant Set Enrichment (VSE) is a fast method to calculate the enrichment of a set of disease-associated variants across functionally annotated genomic regions, consequently highlighting the mechanisms important in the etiology of the disease studied.

**Availability and Implementation:** VSE is implemented as an R package and can easily be implemented in any system with R. See supplementary information for details.

## Introduction

Over 80% of genetic predisposition, namely risk-loci populated by Single Nucleotide Polymorphisms (SNPs), to human diseases identified by Genome Wide Association Studies (GWAS) map to noncoding DNA (Maurano *et al.*, 2012; Cowper-Sal·lari *et al.*, 2012; Schaub *et al.*, 2012). In other words, most disease-associated SNPs do not directly alter coding sequences. Over the last decade, the functional annotation of the coding and noncoding genome across a wide collection of cell and tissue types benefited from the integration of maps of transcriptional activity from both coding and noncoding transcripts, such as miRNA and long-noncoding RNAs (lncRNAs) as well as chromatin-protein binding profiles, inclusive of transcription factors and epigenetics modifications, and open chromatin. This functional annotation provides a unique opportunity to delineate the functional basis of genetic predispositions to disease.

Here, we present a computational method, named Variant Set Enrichment (VSE) that computes the enrichment/depletion of the set of genetic predisposition for a disease of interest over functional genomic annotations. We previously used a VSE-based approach to identify the enrichment of Breast Cancer (BCa) genetic predispositions at enhancers bound by FOXA1 and ESR1 in breast cancer cells (Cowper-Sal·lari *et al.*, 2012). The dependency of VSE on the set of genetic predispositions to a particular disease and on genome-wide functional annotation from a relevant model system renders VSE applicable to the study of any genetically inherited disease.

## Methods

A genetic predisposition (risk-locus) identified by GWAS corresponds to a tagSNP found on the GWAS array and all SNPs missing from the array but known to be in Linkage Disequilibrium (LD) with the tagSNP (Hindorff *et al.*, 2009). The sum of all genetic predispositions to a particular disease, ie: all the tagSNPs and their ldSNPs constitute the Associated Variant Set (AVS) for that disease. The identity of risk-loci is user defined, as the cut-off for LD determination is subjected to study preferences. Occasionally, two or more risk-loci for a particular disease can overlap with one another by a common ldSNP. If the common ldSNP overlaps with a functional genomic annotation of interest, the enrichment score calculated by VSE can be inflated because each risk-locus inclusive of this ldSNP would be counted independently. To correct for this possibility, VSE computes a network of all SNPs in which each SNP is represented as a node and the pairwise LD as an edge. Each cluster in the network represents a disjointed locus, as such, a ldSNP is present only in one locus. VSE then computes the enrichment score of the AVS for each functional genomic annotation of interest in three sequential steps. In the first step, VSE tallies the number of independent risk-loci that overlaps with the functional genomic annotations. Overlapping of a risk-locus is defined as at least one member SNP found within the functional genomic annotation of interest.

This preliminary tallying of AVS may indicate which genomic annotations are functionally related to risk-associated variants, but the overlapping can be affected by size and structure of the AVS. To correct for these biases, VSE, in the second step, computes a null distribution of the overlap tallies that is based on random permutation of AVS. The null AVS is computed by randomly sampling SNPs from a comprehensive pool of tagSNPs present on the GWAS arrays (Illumina Human OmniExpress) and clustering them with their ldSNPs imputed from the 1000 Genome Project Phase III data. When calculating the set of null AVS, VSE makes sure that each set is built in the way that it has identical total number of null loci as the total number of risk-loci in the AVS; and each null locus is matched in size to the corresponding query locus. We defined each null AVS as Matched Random Variant Sets (MRVS).

In the third step, VSE tallies the overlap of MRVS with the functional genomic annotations of interest. This provides the null distribution to calculate for the enrichment/depletion of the AVS across different functional genomic annotations. To make the enrichment analysis comparable across all functional genomic annotations of interest, AVS tally is centered at the median and scaled to the standard deviations of the null distribution. The enrichment score is then defined as the number of standard deviations that the overlapping tally deviates from the null overlapping tally mean. VSE calculates an exact P-value for significance of the enrichment/depletion by fitting a density function to the null distribution derived from the MRVS. The level of significance is corrected for multiple testing using Bonferroni method. The deviation of the null distribution from the normality is tested using Kolmogorov-Smirnov test; and if the distribution deviates, the Box-Cox power transformation is applied on to the null to approach normality.

## Results

The usefulness and impact of VSE is demonstrated by calculating the enrichment of SNPs associated with four cancer types for DNase I Hypersensitivity Sites (DHS) and a set of histone marks profiled genome wide in cancer type relevant cell lines. We compiled 72, 92, 16 and 36 significantly associated SNPs, or tagSNPs, for Prostate Cancer (PRAD), BCa, Lung Cancer (LUAD) and Colorectal Cancer (COAD) respectively from NHGRI catalog (Welter *et al.*, 2014). We computed the LD structure by finding all SNPs in the European population from the 1000 Genome Project that are in LD with the tagSNPs with r2 ≥ 0.8 (The 1000 Genomes Project Consortium, 2012). The DNase-seq and ChIP-seq data for H3K4me1 (enhancer), H3K4me3 (promoter), H3K27ac (enhancer and promoter), H3K36me3 (gene body) and H3K27me3 (repressive region) for each of MCF7 (BCa), LNCaP (PRAD), A549 (LUAD) and HCT-116/Caco-2 (COAD) cell lines are compiled from ENCODE data ENCODE Project Consortium, 2012) and complemented by data from independent studies (Hazelett *et al.*, 2014; Taberlay *et al.*, 2014). Upon performing VSE, the results show that BCa and PRAD AVS are significantly enriched in DHS and regions with H3K27ac mark found in breast and prostate cancer cells, respectively (Fig. 1). On the other hand, SNPs associated with LUAD are enriched in regions with H3K36me3 mark only (Fig. 1). The enrichment of distinct cancer AVS across different functional genomic regions argues for a unique biology affected by genetic predispositions across cancer types.

**Fig. 1:**
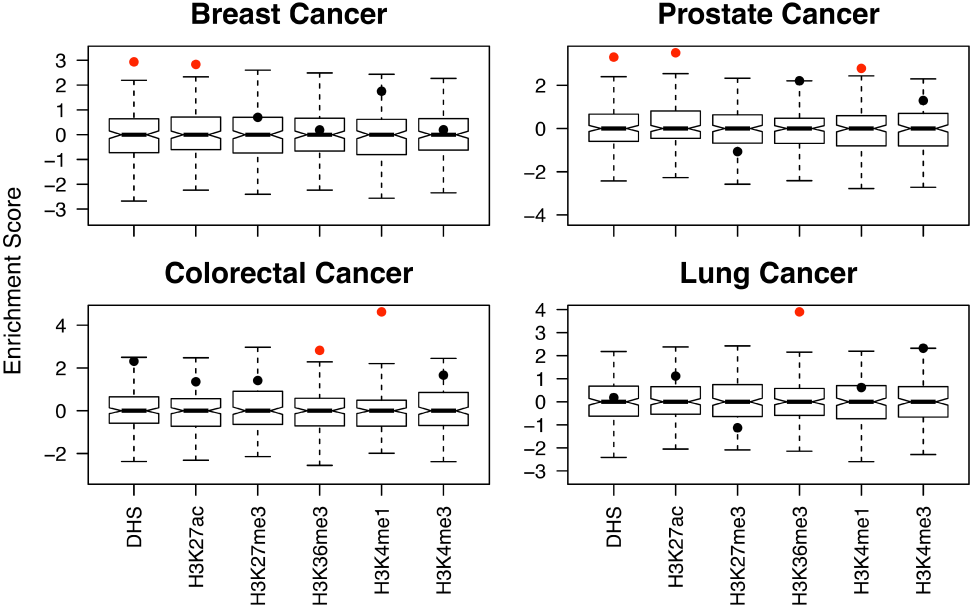
Enrichment of Breast, Prostate, Lung and Colorectal cancer AVS across different genomic maps in cancer-type specific cells. The significantly enriched genome regions (Bonferroni corrected P-value < 0.01) are marked in red. The histone modifications are profiled in MCF7, LNCaP, A549 and HCT-116/Caco-2 for breast, prostate, lung and colorectal cancer, respectively.

## Acknowledgements

We thank Kinjal Desai for helping with setting up the VSE. We acknowledge the ENCODE consortium and the ENCODE production laboratories that generated the data sets provided by the ENCODE Data Coordination Center used in the manuscript.

## Funding

This work was supported by PMCF (to H.H.H. and M.L.), CFI and ORF (CFI32372 to H.H.H.), NSERC (498706 to H.H.H.), WICC and CCS (703800 to H.H.H.), CCSRI (702922 to M.L.), PCC (RS2016-02 to H.H.H; RS2014-04 to M.L.), CIHR (142246 to H.H.H.), NCI at the NIH (R01CA155004 to M.L.; LM009012 and LM010098 to J.H.M.). M.L. holds a young investigator award from the OICR and a new investigator salary award from the CIHR.

Conflict of Interest: none declared.

